# Subcutaneous infusion of neurosecretory protein GL promotes fat accumulation in mice

**DOI:** 10.1101/2021.05.27.446062

**Authors:** Yuki Narimatsu, Keisuke Fukumura, Eiko Iwakoshi-Ukena, Ayaka Mimura, Megumi Furumitsu, Kazuyoshi Ukena

**Author notes:** Correspondence author. E-mail address (K. Ukena). These authors contributed equally to this work (Yuki Narimatsu, Keisuke Fukumura).

## Abstract

We recently identified a novel small secretory protein, neurosecretory protein GL (NPGL), in the vertebrate hypothalamus. We revealed that NPGL is involved in energy homeostasis using intracerebroventricular infusion in rodents. However, the effect of NPGL through peripheral administration remains to be elucidated and may be important for therapeutic use. In this study, we performed subcutaneous infusion of NPGL in mice for 12 days and found that it accelerated fat accumulation in white adipose tissue (WAT) without increasing in body mass gain and food intake. The mass of the testis, liver, kidney, heart, and gastrocnemius muscle remained unchanged. Analysis of mRNA expression by quantitative reverse transcription-polymerase chain reaction showed that proopiomelanocortin was suppressed in the hypothalamus by the infusion of NPGL. We observed a decreasing tendency in serum triglyceride levels due to NPGL, while serum glucose, insulin, leptin, and free fatty acids levels were unchanged. These results suggest that the peripheral administration of NPGL induces fat accumulation in WAT via the hypothalamus.

## 1. Introduction

Recently, the World Health Organization (WHO) reported that 400 million people are obese, and over 1.6 billion adults are overweight worldwide [1]. Obesity is caused by fat accumulation due to excess calorie intake or inadequate energy consumption and has been shown to increase the risk of developing various diseases, including diabetes, hypertension, and hyperlipidemia [2–6]. On the other hand, lack of fat depots also leads to severe diabetes [7]. Thus, there is a strong focus on researching energy homeostasis, including in feeding behavior and lipid metabolism. The regulation of feeding behavior is intricately regulated by neuropeptides in the hypothalamus. Neuropeptide Y (NPY) and agouti-related peptide (AgRP) are orexigenic [8]. It is also known that AgRP competitively inhibits melanocortin receptor 4 (MC4R), the receptor of α-melanocyte-stimulating hormone (α-MSH) derived from proopiomelanocortin (POMC) [9]. In addition, galanin (GAL) controls food intake through GAL receptors, the 5-HT_1A_ receptor, and the adrenergic α-2 receptor [10]. In addition to neuropeptides, peripheral hormones regulate feeding behavior. Insulin, a hormone secreted by the pancreas, suppresses feeding behavior, mediated by cocaine- and amphetamine-regulated transcript (CART) in the hypothalamus [11]. Orexigenic/anorexigenic factors and their signaling pathways regulate not only feeding behavior but also lipid metabolism. Central MC4R signaling suppresses lipogenesis in the peripheral adipose tissue by activating the sympathetic nervous system [12]. Insulin promotes lipogenesis by inhibiting lipolysis [13]. Moreover, ghrelin, a hormone secreted by the stomach, stimulates fat accumulation by reducing fat utilization [14]. Additionally, leptin plays an important role in the regulation of lipolysis [15]. The mechanism of lipid metabolism is not fully understood, and further research is required to regulate fat mass in various tissues.

We recently identified a novel cDNA-deduced precursor of a small hypothalamic secretory protein as a regulator of energy homeostasis in chickens, rats, mice, and humans [16–18]. The precursor protein includes a signal peptide sequence, a glycine amidation signal, and a cleavage site. A mature small protein was termed neurosecretory protein GL (NPGL) because it contains 80 amino acid residues with a specific sequence of Gly-Leu-NH_2_ in the C-terminus [16]. Similar to NPGL, we have also identified the neurosecretory protein GM (NPGM), a paralogous protein of NPGL in vertebrates [16]. We found that chronic intracerebroventricular (i.c.v.) infusion of NPGL induced feeding behavior and fat accumulation in rats and mice [17,19]. In addition, our previous study revealed that NPGL was specifically expressed in the hypothalamus of mice [18]. The effect of subcutaneous NPGL infusion is unknown, and understanding the effect of NPGL on the periphery is necessary for its potential therapeutic use.

In this study, we subcutaneously infused NPGL and investigated its peripheral impact on body composition, feeding behavior, and serum levels of biochemical parameters in mice. Furthermore, mRNA expression of lipid metabolism-related factors in white adipose tissue (WAT) and neuropeptides in the hypothalamus was examined using quantitative reverse transcription-polymerase chain reaction (qRT-PCR).

## 2. Materials and methods

### 2.1. Animals

Male C57BL/6J mice (7 weeks old) were purchased from Nihon SLC (Hamamatsu, Japan) and housed under standard conditions (25 ± 1 °C under a 12-hr light/dark cycle) with *ad libitum* access to water and a high-calorie diet (32% of calories from fat/20% of calories from sucrose, D14050401; Research Diets, New Brunswick, NJ, USA). All animal experiments were performed according to the Guide for the Care and Use of Laboratory Animals prepared by Hiroshima University (Higashi-Hiroshima, Japan), and these procedures were approved by the Institutional Animal Care and Use Committee of Hiroshima University (permit number: C19-8).

### 2.2. Production of AGIA-NPGL

NPGL is a hydrophobic protein and the AGIA tag, a hydrophilic affinity tag [20], was labeled at the N-terminus of NPGL. AGIA-NPGL, containing 89 amino acid residues, was synthesized by microwave-assisted solid-phase peptide synthesis using an automated peptide synthesizer (Initiator+ Alstra; Biotage, Uppsala, Sweden) as previously described [21].

### 2.3. Subcutaneous infusion of NPGL

For the 12-day chronic subcutaneous infusion of NPGL, we used an Alzet mini-osmotic pump (model 2001, delivery rate 1.0 μL/h; DURECT Corporation, Cupertino, CA, USA). NPGL (24 nmol/day) was dissolved in 30% propylene glycol and adjusted to pH 8.0 with NaOH. For control animals, a vehicle solution was used. The dose of NPGL was determined based on our previous study that subcutaneous administration of 24 nmol/day NPGM was effective on body mass gain, while that of low dose (2.4 nmol/day) did not show any effects in chicks [22]. Prior to subcutaneous pump implantation onto the back of mice, pumps were filled with vehicle or NPGL solution and soaked in 0.9% NaCl at 37°C overnight. The pumps were replaced with new ones 6 days after their first implantation. We confirmed correct infusion by examining the remaining solution in the pump after its removal.

### 2.4. Measurement of body mass, food intake, and body composition

Body mass and food intake were recorded every day. After 12 days of the first implantation of the osmotic pumps, mice were immediately decapitated between 13:00–15:00. The adipose tissues, organs, skeletal muscles, and hypothalamus were collected, weighed, and frozen in liquid nitrogen. Blood was collected at the same time when mice were sacrificed.

### 2.5. Serum biochemical analysis

Serum levels of glucose, lipids, and hormones were measured. The GLUCOCARD G+ (Arkray, Kyoto, Japan) meter was used to measure glucose content. The Rebis Insulin-mouse T ELISA kit (Shibayagi, Gunma, Japan) was used to measure insulin levels. The Leptin ELISA kit (Morinaga Institute of Biological Science, Yokohama, Japan) was used to measure leptin levels. The NEFA C-Test Wako (Wako Pure Chemical Industries, Osaka, Japan) was used to measure free fatty acid levels. Triglyceride E-Test Wako (Wako Pure Chemical Industries) was used to measure triglyceride levels.

### 2.6. qRT-PCR

The inguinal WAT and the hypothalamus were dissected from mice and snap frozen in liquid nitrogen for RNA processing after NPGL infusion. Total RNA was extracted using QIAzol lysis reagent for WAT (QIAGEN, Venlo, Netherlands) or TRIzol reagent for the hypothalamus (Life Technologies, Carlsbad, CA, USA) according to the manufacturer’s instructions. First-strand cDNA was synthesized from total RNA using a ReverTra Ace kit (TOYOBO, Osaka, Japan). We analyzed the following factors in WAT: acetyl-CoA carboxylase (ACC), fatty acid synthase (FAS), stearoyl-CoA desaturase 1 (SCD1), and glycerol-3-phosphate acyltransferase 1 (GPAT1) as lipogenic enzymes; carbohydrate-responsive element-binding protein α (ChREBPα) as a lipogenic transcription factor; carnitine palmitoyltransferase 1a (CPT1a), adipose triglyceride lipase (ATGL), and hormone-sensitive lipase (HSL) as lipolytic enzymes; glyceraldehyde-3-phosphate dehydrogenase (GAPDH) as a carbohydrate metabolism enzyme; solute carrier family 2 member 4 (SLC2A4) as a glucose transporter; cluster of differentiation 36 (CD36) as a fatty acid transporter; peroxisome proliferator-activated receptor (PPAR) α and γ as lipid-activated transcription factors; and tumor necrosis factor α (TNFα) as an inflammatory cytokine. We analyzed the following factors in the hypothalamus: NPGL, NPGM, NPY, and AgRP as orexigenic neuropeptides; POMC as an anorexigenic neuropeptide; GAL as a neuropeptide that enhances fat intake. The abbreviations for genes and primer sequences used in this study are listed in Table 1 and Table 2, respectively. PCR amplifications were conducted with THUNDERBIRD SYBR qPCR Mix (TOYOBO) using the following conditions: 95 °C for 20 s, followed by 40 cycles of 95 °C for 3 s, and 60 °C for 30 s. The PCR products in each cycle were monitored using Bio-Rad CFX Connect (Bio-Rad Laboratories, Hercules, CA, USA). Relative quantification of each gene was determined by the 2^−∆∆Ct^ method using ribosomal protein S18 (*Rps18*) for WAT or β-actin (*Actb*) for the hypothalamus as internal controls [23].

**Table 1.** Abbreviation

| Gene | Name |
| --- | --- |
| *Acc* | Acetyl-CoA carboxylase |
| *Fas* | Fatty acid synthase |
| *Scd1* | Stearoyl-CoA desaturase 1 |
| *Gpat1* | Glycerol-3-phosphate acyltransferase 1 |
| *Chrebpα* | Carbohydrate-responsive element-binding protein $\alpha$ |
| *Cpt1a* | Carnitine palmitoyl transferase 1a |
| *Atgl* | Adipose triglyceride lipase |
| *Hsl* | Hormone-sensitive lipase |
| *Gapdh* | Glyceraldehyde 3-phosphate dehydrogenase |
| *Slc2a4* | Solute carrier family 2 member 4 |
| *Cd36* | Cluster of differentiation 36 |
| *Ppara* | Peroxisome proliferator-activated receptor $\alpha$ |
| *Pparγ* | Peroxisome proliferator-activated receptor $\gamma$ |
| *Tnfa* | Tumor necrosis factor $\alpha$ |
| *Rps18* | Ribosomal protein S18 |
| *Npgl* | Neurosecretory protein GL |
| *Npgm* | Neurosecretory protein GM |
| *Npy* | Neuropeptide Y |
| *AgRP* | Agouti-related peptide |
| *Pomc* | Proopiomelanocortin |
| *Gal* | Galanin |
| *Actb* | β-Actin |

**Table 2.**
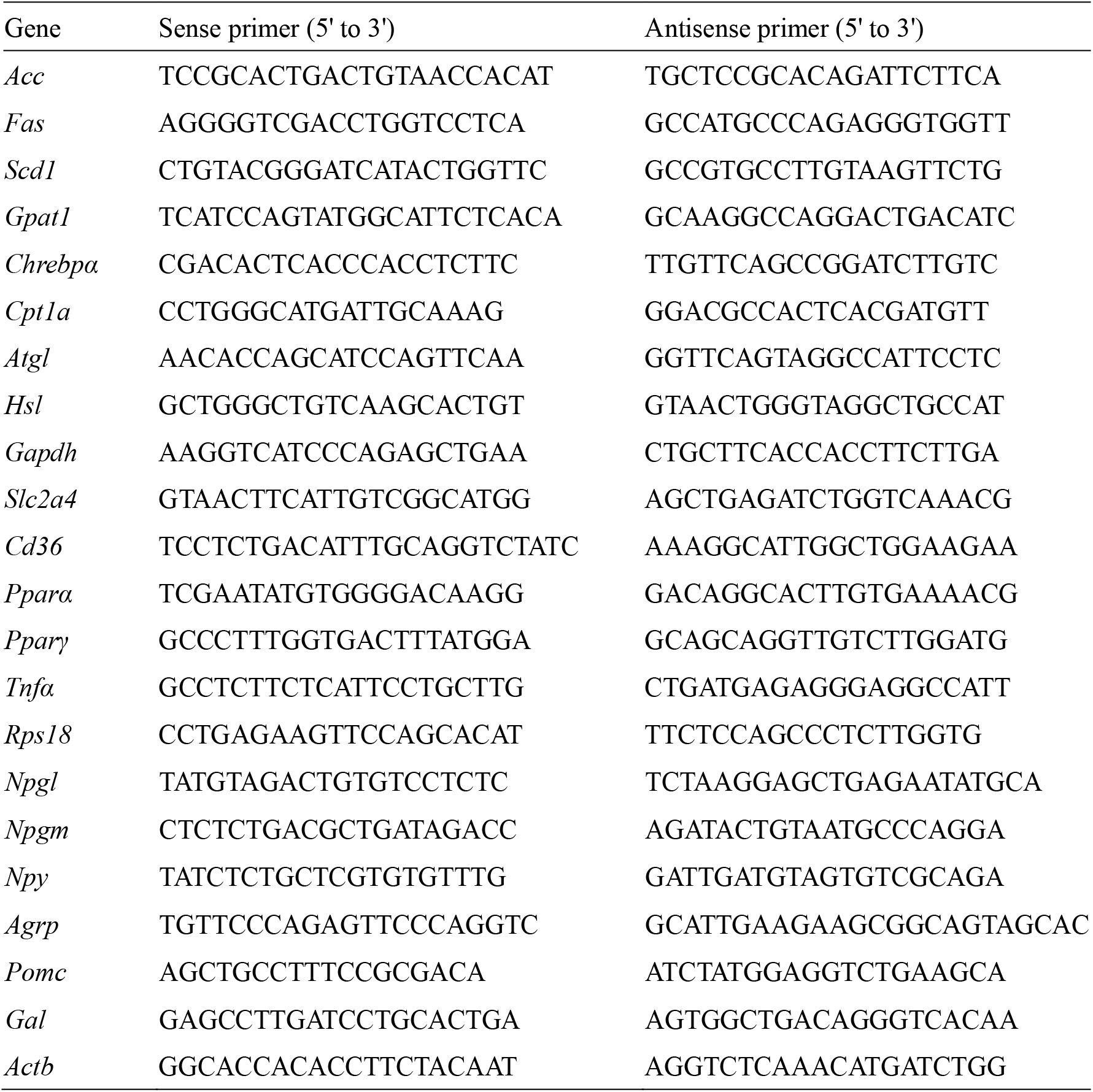
Sequences of oligonucleotide primers for qRT-PCR

### 2.7. Statistical analysis

Group differences between NPGL and vehicle-treated animals were assessed using Student’s *t-*tests for tissue mass and mRNA expression. One-way analysis of variance (ANOVA) followed by Bonferroni’s test, as appropriate, was used for analyzing group differences regarding body mass and food intake. *P* values of <0.05 were considered statistically significant.

## 3. Results

### 3.1. Effects of subcutaneous infusion of NPGL on body mass gain, food intake, and body composition

Chronic subcutaneous administration of NPGL (24 nmol/day) was conducted using osmotic pumps in mice fed a high-calorie diet. Although NPGL slightly increased body mass gain and cumulative food intake without significant differences (Fig. 1A and B), it significantly increased the mass of inguinal WAT and perirenal WAT (Fig. 2A). In contrast, the mass of epididymal WAT, retroperitoneal WAT (Fig. 2A), and interscapular BAT (Fig. 2B) did not increase. Moreover, the mass of the testis, liver, kidney, heart (Fig. 2C), and gastrocnemius muscle (Fig. 2D) remained unchanged.

**Fig. 1.**
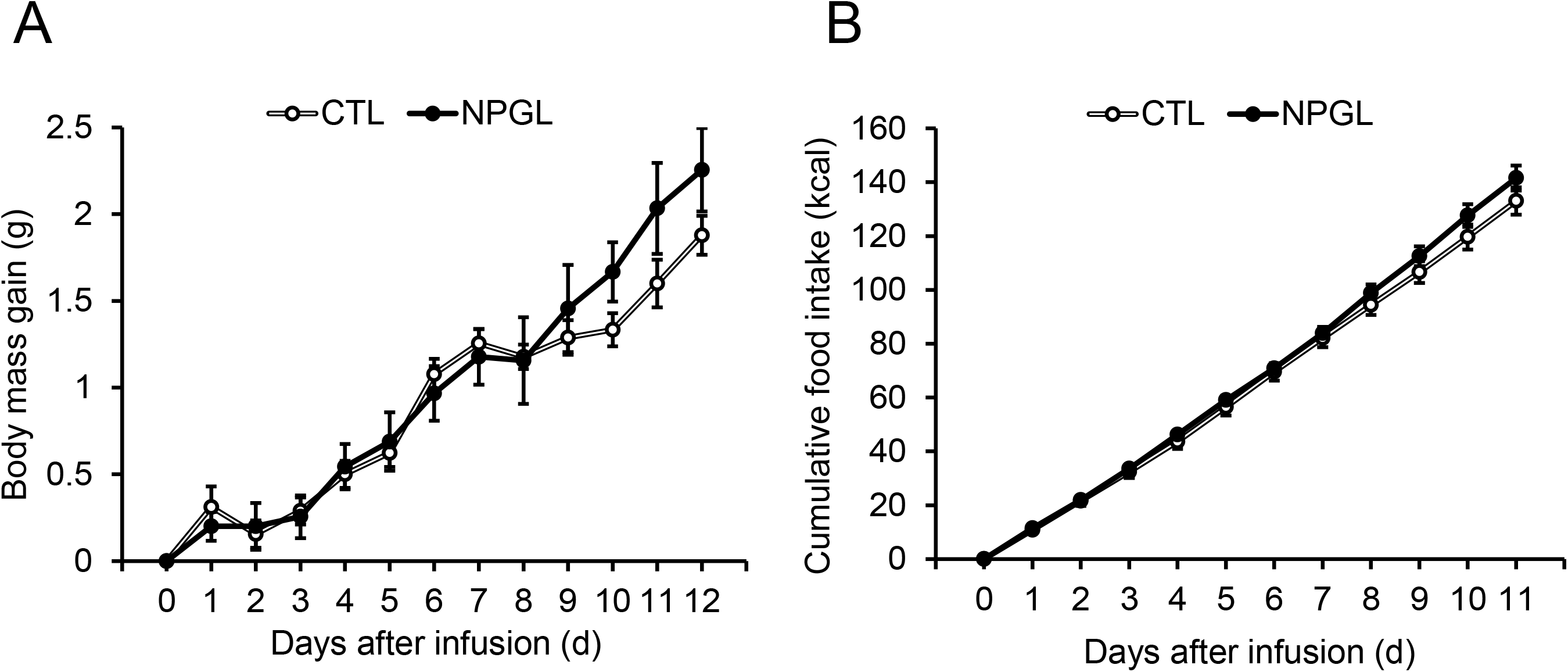
The effects of subcutaneous infusion of NPGL on body mass gain and food intake. (A) Body mass gain. (B) Cumulative food intake. Each value represents the mean ± standard error of the mean (n=9/group). CTL, control animal group treated with a vehicle solution; NPGL, experimental animal group treated with neurosecretory protein GL.

**Fig. 2.**
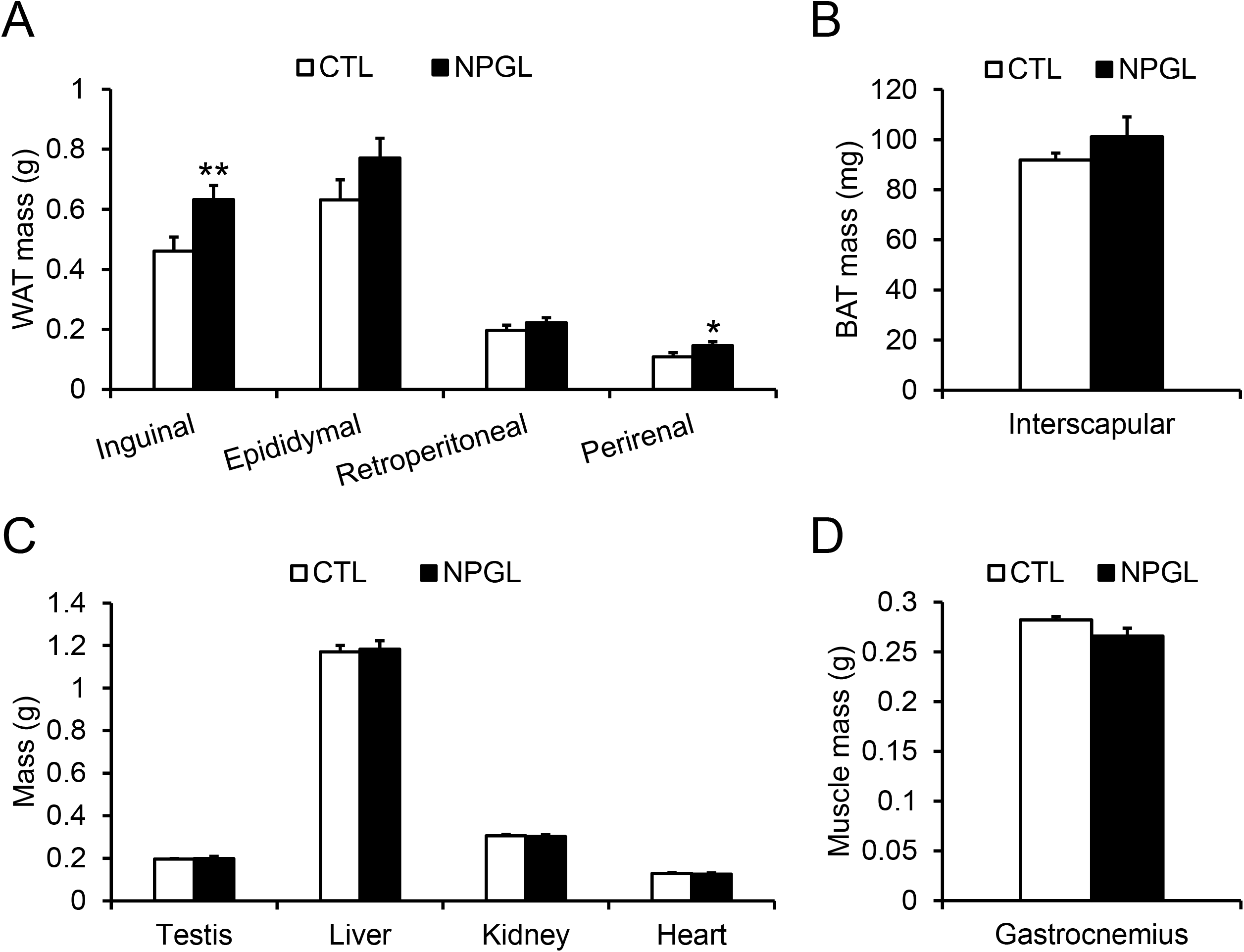
The effects of subcutaneous infusion of NPGL on body composition. (A) Mass of inguinal, epididymal, retroperitoneal, and perirenal white adipose tissue (WAT). (B) Mass of interscapular brown adipose tissue (BAT). (C) Mass of testis, liver, kidney, and heart. (D) Mass of gastrocnemius muscle. Each value represents the mean ± standard error of the mean (n=9/group). \**P*<0.05, \*\**P*<0.01. CTL, control animal group treated with a vehicle solution; NPGL, experimental animal group treated with neurosecretory protein GL.

### 3.2. Effects of subcutaneous infusion of NPGL on mRNA expression of lipid metabolism-related genes

After observing subcutaneous infusion of NPGL-stimulated fat accumulation in inguinal WAT and perirenal WAT, we analyzed mRNA expression of genes involved in lipid metabolism by qRT-PCR. qRT-PCR showed that subcutaneous infusion of NPGL induced no change in the mRNA expression of these genes in inguinal WAT (Fig. S1). To examine the effect of NPGL on the hypothalamus after subcutaneous infusion, we measured the mRNA expression for several neuropeptides and found that POMC mRNA was downregulated (Fig. 3).

**Fig. 3.**
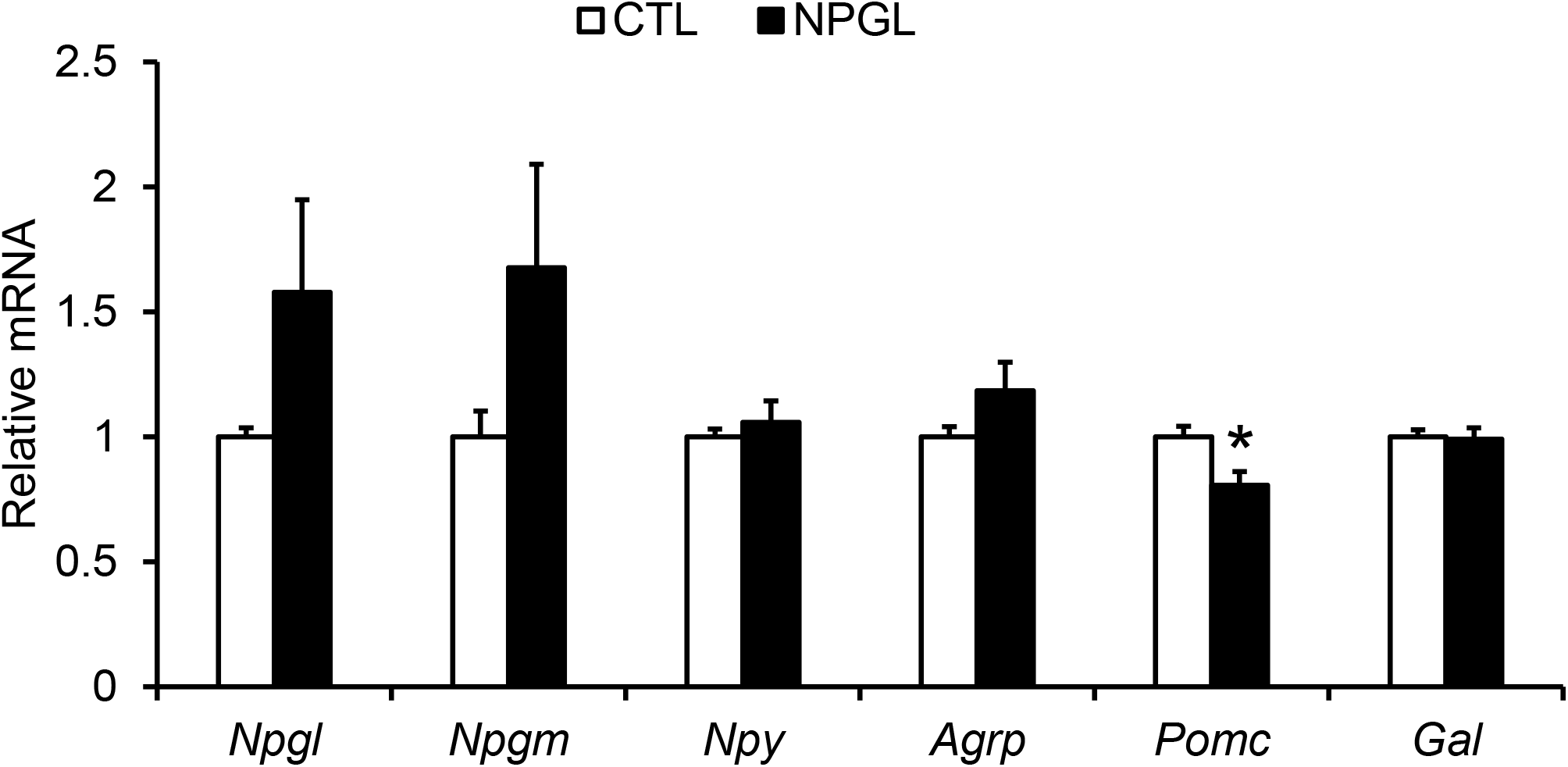
The effects of subcutaneous infusion of NPGL on mRNA gene expression. mRNA expression levels of neuropeptides in the hypothalamus. Each value represents the mean ± standard error of the mean (n=9/group). \**P*<0.05. CTL, control animal group treated with a vehicle solution; NPGL, experimental animal group treated with neurosecretory protein GL.

### 3.3. Effects of subcutaneous infusion of NPGL on serum parameters

Because the mass of WAT increased by infusion of NPGL, we analyzed serum levels of glucose, hormones, and lipids. Although the serum level of triglycerides tended to decrease with NPGL (Fig. 4D), the levels of glucose, insulin, leptin, and free fatty acids remained unchanged (Fig. 4A, B, C, E).

**Fig. 4.**
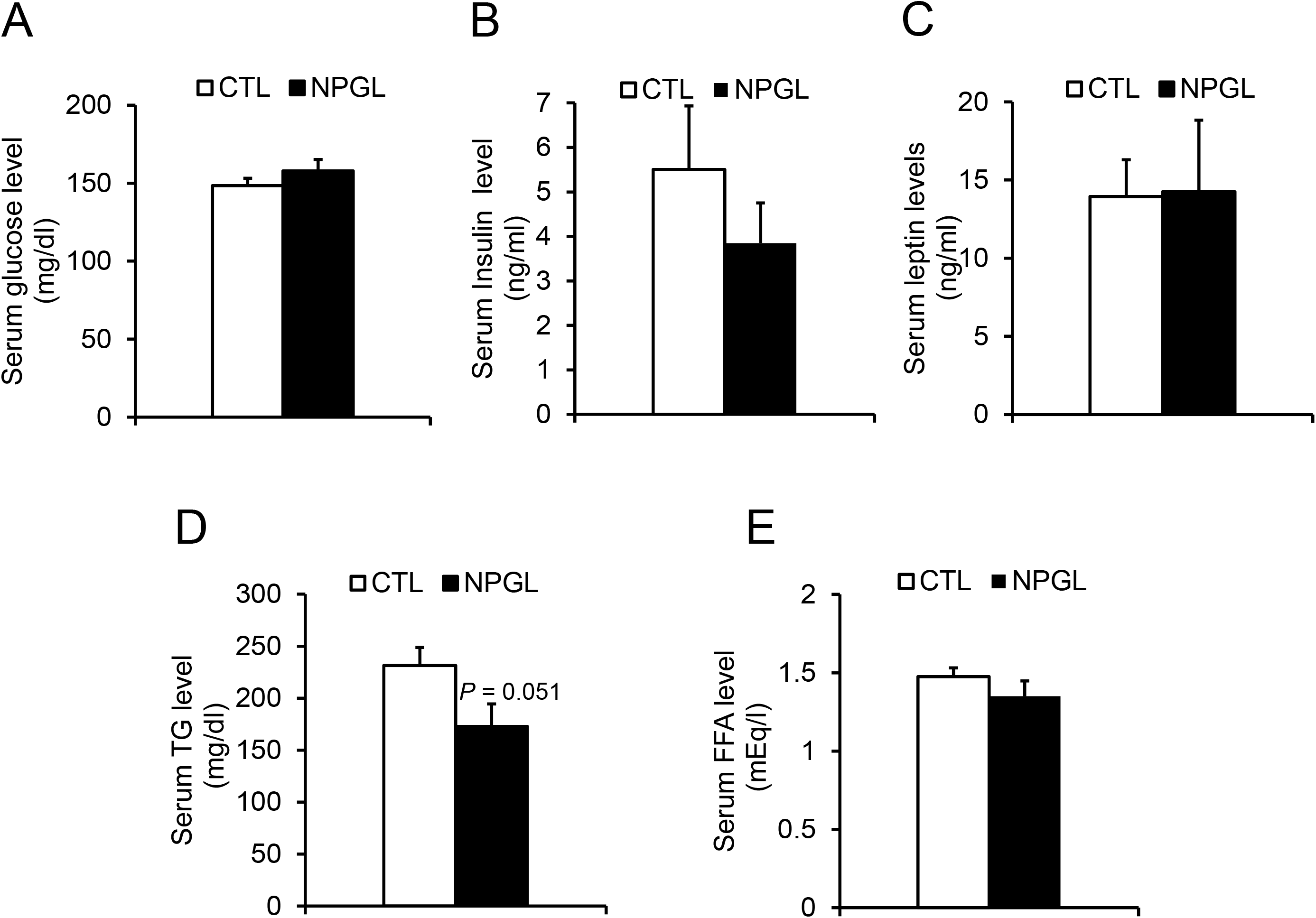
Serum parameters during subcutaneous infusion of NPGL. (A) Serum glucose. (B) Serum insulin. (C) Serum leptin. (D) Serum triglyceride (TG). (E) Serum free fatty acids (FFA). Each value represents the mean ± standard error of the mean (n=9/group). CTL, control animal group treated with a vehicle solution; NPGL, experimental animal group treated with neurosecretory protein GL.

## 4. Discussion

We have reported that NPGL is involved in fat accumulation using chronic i.c.v. infusion and/or overexpression of the precursor gene in chickens, rats, and mice [16,17,19,24]. However, the peripheral effect of NPGL in mammals has not been elucidated. In this study, we demonstrated that subcutaneous infusion of NPGL stimulated fat accumulation in WAT, corresponding to our previous study showing that i.c.v. infusion of NPGL enlarged adipocytes in WAT [17,19]. In contrast, body mass gain, food intake, and blood parameters seemed to be hardly affected by NPGL. These data suggest that NPGL stimulates fat accumulation in WAT through peripheral infusion, as well as through central action.

The present study also showed that mRNA expression of POMC, the precursor protein of anorexigenic α-MSH, was significantly decreased in the hypothalamus after peripheral administration of NPGL. Morphological analysis revealed that NPGL neurons were localized in the lateroposterior part of the arcuate nucleus (Arc), and their fibers projected to the rostral region of the Arc [18]. The Arc is located close to the median eminence (ME), which is deficient in the blood-brain barrier (BBB) [25]. Indeed, it is well known that some peripheral hormones act on the neurons in the Arc [26–28]. For instance, peripheral leptin decreases lipid deposition in adipose tissues through regulation of POMC neurons in the Arc [29]. In addition, we have found that NPGL fibers innervate to POMC neurons and i.c.v. infusion of NPGL decreased energy expenditure in mice [18,19]. Thus, it is possible that subcutaneous infusion of NPGL may decrease energy expenditure by inhibiting POMC neurons in the Arc through the ME, resulting in fat accumulation in WAT. To date, the receptor for NPGL has not been identified. Further work is needed to characterize the NPGL receptor and its binding site in the brain, including POMC neurons.

NPGL stimulated fat accumulation in WAT, while a decrease in serum triglyceride levels was observed (Figs. 2A, 4D). It has been reported that serum triglyceride levels are regulated by a neuronal relay of liver-brain-adipose tissue [30]. When the liver detects circulating amino acids in an energy-rich state, it transmits a signal to the brain through the vagal afferent pathway [30]. Subsequently, the brain activates the sympathetic nervous system to suppress mRNA expression of lipoprotein lipase (LPL) in WAT [30]. Finally, serum triglyceride levels are increased by inhibiting the degradation of triglycerides [30]. In this way, serum triglyceride is increased by activation of sympathetic nerve from the brain to the adipose tissue [30]. Indeed, our recent work suggests that NPGL suppresses sympathetic nerve activity promoting lipogenesis in rats (unpublished data). Therefore, the decrease in serum triglyceride levels may be due to the suppression of sympathetic nerve activity by NPGL. Future studies are needed to investigate in detail the impact of NPGL on sympathetic regulation in mice.

While we expanded the potential of NPGL as a therapeutic agent for metabolic diseases, there were several limitations to the present study. First, we could not validate the effect of subcutaneous infusion of NPGL on lipid metabolism in a dose-dependent manner in the present study, while we previously reported the dose-dependent effects of NPGM in chicks [22]. Further study is required to address the dose-dependent effect of NPGL in mice. Second, the present study revealed that NPGL stimulated fat accumulation without changing the mRNA expression involved in lipid metabolism. Further analysis of the effects of NPGL on lipid-metabolic enzymes at translational and phosphorylational levels will help understand the molecular mechanisms underlying fat accumulation induced by NPGL. Third, we could not address the effects of dietary nutrition on NPGL activity in the present study, although subcutaneous infusion of NPGL induced fat accumulation in mice fed a high-calorie diet. Indeed, several previous studies implied that NPGL exhibited multiple effects on energy metabolisms [31,32]. Therefore, future studies using various diets, including a normal diet, will open up a new avenue for the therapeutic use of subcutaneous infusion of NPGL in different nutritional conditions.

## 5. Conclusion

Besides i.c.v. injection, subcutaneous infusion of NPGL also promotes fat accumulation in WAT in mice. The infusion of NPGL did not increase body mass and food intake. Despite fat accumulation, we observed a tendency for serum triglyceride levels to decrease; however, serum levels of glucose, insulin, leptin, and free fatty acids did not change. Furthermore, the peripheral administration of NPGL decreased the mRNA expression of POMC, one of precursors of catabolic factors, in the hypothalamus, while that of NPGL did not affect the mRNA expression of lipid-metabolic factors in WAT. These results suggest that the peripheral infusion of NPGL may inhibit the activity of POMC neurons through the BBB and induce fat accumulation in WAT. This is the first report describing a biological action of peripheral administration of NPGL in mammals.

## Acknowledgements

We are grateful to Mr. Atsuki Kadota and Ms. Mana Naito for the experiment support. This work was supported by JSPS KAKENHI Grant (JP18K19743, JP19H03258, JP20K21760, and JP20H03296 to K.U., JP19K06768 to E.I.-U., and JP20K22741 to K.F.), the Takeda Science Foundation (K.U.), the Uehara Memorial Foundation (K.U.), the ONO Medical Research Foundation (K.U.), and the Electric Technology Research Foundation of Chugoku (K.U.).

