## Supplementary Figure for "Subcutaneous infusion of neurosecretory protein GL promotes fat accumulation in mice"

Fig. S1

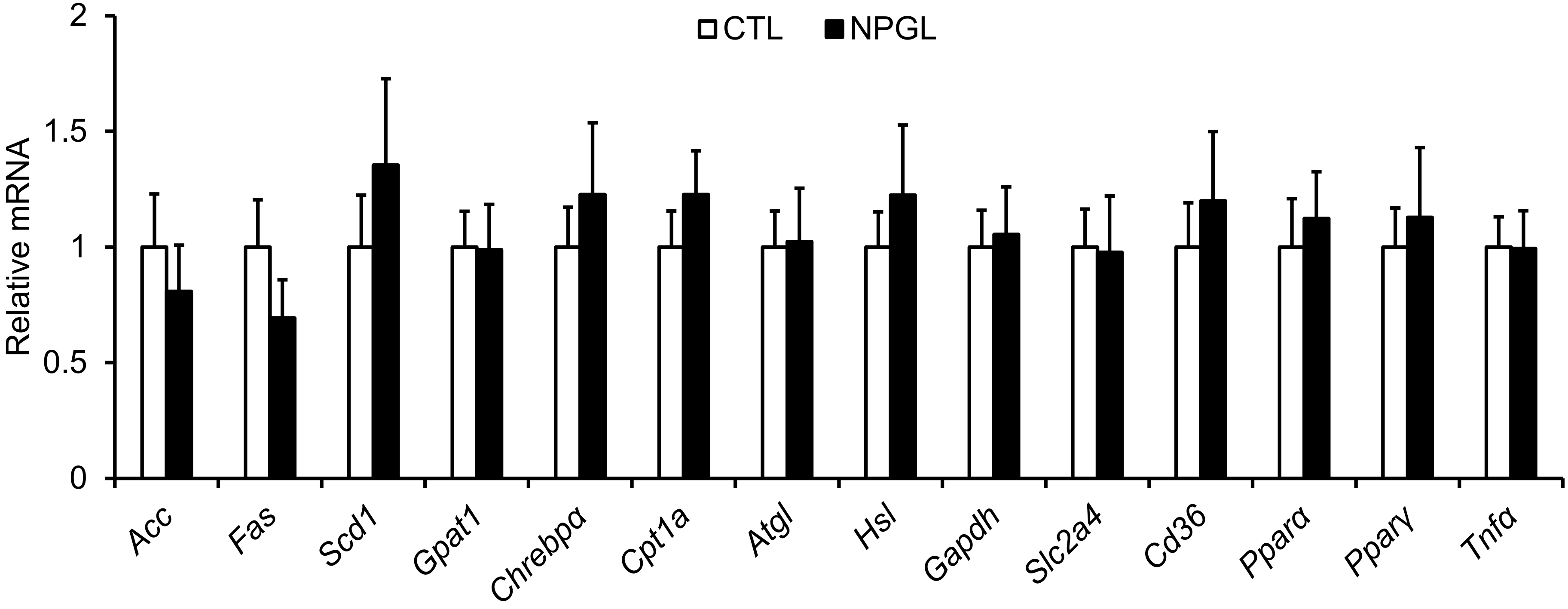


**Supplementary Fig. S1.**
The effects of subcutaneous infusion of NPGL on mRNA gene expression. mRNA expression levels related to lipid metabolism for inguinal WAT. Each value represents the mean ± standard error of the mean (n=9/group). CTL, control animal group treated with a vehicle solution; NPGL, experimental animal group treated with neurosecretory protein GL.
